# Innate building blocks underly socially learned call sequences

**DOI:** 10.64898/2026.06.21.733568

**Authors:** Stephanie L. Mason, Sarah L. Walsh, Amanda R. Ridley

## Abstract

Recent evidence of extensive call sequence use in non-human primates has led to the theory that syntax evolved to mitigate the constraints of their genetically fixed repertoires, before vocal production learning later emerged in humans. However, evidence of similarly extensive sequence repertoires in an open-ended vocal production learner—the Western Australian magpie (*Gymnorhinatibicendorsalis*)—offers a unique opportunity to explore potential alternative pathways to syntactic communication. Our previous work revealed fledgling magpies learn group-specific repertoires of structured call sequences from their social contacts, with more sociable individuals acquiring larger repertoires earlier in development. Notably however, the individual vocal ‘segments’ that combine to form their calls and call sequences were shared across groups and emerged as early as the first week post-fledging—suggesting the underlying vocal elements may not be learned. Here we utilised acoustic neighbourhood-based dimensionality reduction to compare clustering patterns of vocal segments across magpie fledgling developmental stages, and between fledglings and adults. We found no evidence of acoustic development over time, and no significant distinction between fledgling and adult productions of the same vocal segments. The same coarticulatory effects—where a vocal element is produced differently when combined with another—and geographic variation established previously in adults were supported in fledglings too. These findings support that the vocal building blocks underpinning magpie call sequences are innate, suggesting usage learning better explains how fledglings learn to combine calls. In a species capable of open-ended production learning, this suggests learning to combine existing signals may be more adaptive than productively learning new ones. Rather than evolving solely to compensate for genetically fixed repertoires, syntax may have evolved as a flexible, convergent solution to the various challenges of expanding communicative capacity—whether due to genetic constraints, cognitive limitations or the cost of establishing new meaning in novel signals.

## Introduction

Syntax is the ability to combine meaningful signals (e.g., words) that are themselves made up of ordered, meaningless sounds (e.g., phonemes), into meaningful, ordered sequences (e.g., sentences). This capacity—critical to our impressive communicative flexibility and generativity—was long considered unique to human language (Hurford, 2012; Marler, 1998). While structural similarities to syntax have long been studied in how syllables are combined within animal song (Berwick et al., 2011; Doupe & Kuhl, 1999; Garland et al., 2017; Hurford, 2012; Katahira et al., 2011; Marler & Pickert, 1984; Menyhart et al., 2015; Payne & McVay, 1971; Sainburg et al., 2019; Soma et al., 2009; Tchernichovski et al., 2001), the lack of discrete, semantic meaning in song limits comparison to linguistic syntax, for which meaning is crucial (Berwick et al., 2011; Bolhuis et al., 2010; Engesser & Townsend, 2019). However, in recent years, evidence of non-human animals (hereafter animals) combining discrete, meaningful calls has accumulated (Berthet et al., 2019, 2025; Bosshard et al., 2024; Engesser et al., 2024; Engesser & Townsend, 2019; Girard-Buttoz, Zaccarella, et al., 2022; Hedwig & Kohlberg, 2024; Selbmann et al., 2023; Suzuki et al., 2016, 2017; Walsh et al., 2023), providing the opportunity for investigating the evolutionary origins of syntactic communication both as a homologous trait in various primates (Berthet et al., 2025; Bortolato et al., 2023; Bosshard et al., 2024; Girard-Buttoz, Zaccarella, et al., 2022), and an analogous trait in more distantly related species (e.g., songbirds, Engesser et al., 2016; Suzuki et al., 2016; Walsh et al., 2023; mongooses, Collier et al., 2017; canids, Déaux et al., 2016; and elephants, Hedwig & Kohlberg, 2024).

To date, most studies into semantic combinatoriality have looked primarily at single vocalisations (Engesser et al., 2016, 2019; Leroux et al., 2021; Suzuki et al., 2016) and provided evidence of relatively simple, two-call sequences (i.e., bigrams, Engesser et al., 2016; Leroux et al., 2021; Ouattara et al., 2009; Suzuki et al., 2016). Without taking a whole-repertoire approach, it remains unclear whether there is a genuine discontinuity between the flexibility and diversity of syntax in language and other animal systems, or whether we simply have yet to uncover the true extent of animal syntax. To close this gap, several recent studies have taken a whole-repertoire approach to studying combinatoriality and in doing so, have uncovered the most extensive examples of call sequence usage to date (Bosshard et al., 2024; Girard-Buttoz, Zaccarella, et al., 2022; Mason, King, et al., 2026; Walsh et al., 2023). Chimpanzees (*Pan troglodytes*) have been found to produce 390 unique sequences of up to 10 calls long, with 15% of the total vocal output measured in one study comprising sequences of 3 -10 calls (Girard-Buttoz, Zaccarella, et al., 2022). Common marmosets (*Callithrix jacchus*) have similarly been found to produce sequences of 2-9 calls where calls are arranged following predictable second order rules—where calls are predicted by both preceding calls together (Bosshard et al., 2024). Evidence of such complex sequences in primates both closely and distantly related to humans, suggest the capacity for syntax may have evolved in a common ancestor some 45 million years ago (Leroux & Townsend, 2020). Non-human primates seem to lack the capacity for vocal production learning—the ability to learn new vocalisations from heard models (Fischer & Hammerschmidt, 2020; Janik & Knörnschild, 2021). As such, one theory for the evolution of syntax is that it emerged to mitigate the communicative constraints of genetically fixed repertoires, with humans evolving the capacity for vocal production learning to increase communicative complexity even further following our divergence from other primates (Bortolato et al., 2023; Nowak et al., 2000). Western Australian magpies, like humans, are open-ended vocal production learners—not only can they learn to produce novel vocalisations, but they can do so across their entire lifespan (Kaplan, 1999; Suthers et al., 2011). Recent evidence has shown magpies combine discrete calls to produce over 100 unique call sequences up to 15 calls long, with predictable ordering rules both for how segments (akin to phonemes) are combined into calls and how calls are combined into sequences—the first evidence of multi-level structure in any semantic animal call system (Mason, King, et al., 2026; Mason, Walsh, et al., 2026; Walsh et al., 2023). The co-occurrence of open-ended vocal production learning and combinatoriality, particularly in such a distant relative, provides the opportunity to explore alternative theories for why syntax may have evolved, independent of phylogenetic relatedness.

To date very few studies have looked at how call sequences develop within species, limiting our capacity to understand the drivers of this capacity within the species in which it is found, and by extension, its broader evolutionary drivers. Two of the three ontogenetic studies into call sequence development of which we are aware have been cross-sectional studies or single observations of primates, preventing investigation into the potential interplay between vocal production learning and call combining and limiting investigation of both within- and between-individual variation. While one two-call chimpanzee sequence has been shown to vary in order between two populations (A-B versus B-A; Girard-Buttoz, Bortolato, et al., 2022), magpie sequence repertoires are highly variable between groups, with larger groups producing a greater diversity of sequences (Walsh et al., 2024). Mason, King et al. (2026) performed the first longitudinal study of call sequence development and used individual-level measures of vocal and social complexity to investigate what drives sequence variation. Fledglings were found to learn their call sequences from their social groups, with more sociable fledglings learning more sequences earlier in development—the first evidence of an animal learning to combine meaningful calls into sequences (Mason, King, et al., 2026). Interestingly, all magpie sequences are comprised of the same underlying calls, some of which emerge as early as the first week out of the nest, and calls show little increase in diversity with age (Mason, King, et al., 2026). Additionally, while the diversity and rate of development of sequences is driven by individual sociability, the discrete calls that comprise them develop similarly in all fledglings with no effect of sociability (Mason, King, et al., 2026). These findings suggest that the calls themselves may be innate, and instead only the way in which they are combined into sequences is learned from the social group. Confirming this would help us better understand the mechanisms underlying magpie call sequencing, and how and why this capacity may have emerged within theoretically unconstrained repertoires.

One approach to this, is to determine whether calls undergo acoustic development during ontogeny. While acoustic development is not evidence of vocal production learning (see Fitch & Hauser, 1995; Hammerschmidt et al., 2001; Seyfarth & Cheney, 1997; Snowdon et al., 1997, for evidence of acoustic development in innate primate vocalisations), it is a prerequisite for it (Egnor & Hauser, 2004). As such, calls that show no acoustic development over the ontogenetic period are unlikely to be learned. Determining whether magpie calls develop acoustically over time or are adult-like at first production could inform our understanding of their developmental basis. Here, we analyse the acoustic structure of vocalisations captured in our longitudinal focal study (Mason, King, et al., 2026), utilising an emerging dimensionality reduction technique for classifying vocalisations: uniform manifold approximation and projection (UMAP: Sainburg et al., 2020; Thomas et al., 2022). While dimensionality reduction has been used extensively to study vocalisations, most methods use a single acoustic parameter (e.g. fundamental frequency or mean spectral entropy) meaning the results can vary greatly depending on the acoustic parameter chosen (Thomas et al., 2022). UMAP analyses spectrograms holistically, retaining a more objective measure of all their acoustic properties (Sainburg et al., 2020; Thomas et al., 2022). Thus far, UMAP has been utilised to identify unique vocal elements (such as in magpies; Walsh et al., 2023), as well as individual and sex differences in various species’ repertoires (meerkats *Suricata suricatta*, Thomas et al., 2022; plains zebras *Equus quagga*, Xie et al., 2024; and rooks *Corvus frugilegus*, Martin et al., 2024). Using magpie vocal labels established in Walsh et al. (2023) we analyse the acoustic structure of fledgling vocalisations across the ontogenetic period to determine (i) if fledgling and adult productions of the same vocal element differ significantly and (ii) whether fledgling vocalisations cluster differently at three developmental stages (1-10 weeks, 11-20 weeks and 21-30 weeks post-fledging).

## Methods

### Study species and population

Western Australian magpies are passerines that live in stable, cooperatively breeding groups of 2 to 12 adult individuals (Ashton et al., 2018; Pike et al., 2019). Fledgling magpies are fed by adults for approximately 100 days (Ashton et al., 2018), before they become nutritionally independent. Magpies become sexually mature between 1 to 3 years—when they attain their adult plumage and begin nesting—and live for up to 30 years (Rowley et al., 2022). Magpies’ capacity for open-ended vocal production learning is evidenced by their novel mimicry of other animal calls and anthropogenic sounds that they produce throughout life (Kaplan, 1999; Suthers et al., 2011). Magpies also produce song which can range from solo warbles to group choruses (Brown & Farabaugh, 1991; Kaplan, 1999), and non-song vocalisations in the form of discrete calls and call sequences (Dutour et al., 2020, 2023; Walsh, 2024; Walsh et al., 2019, 2023). Magpies produce four distinct vocal segments: *noisy lines* (*NL*), *down sweeps* (*DS*), *long highs* (*LH*) and *short highs* (*SH*) (Figure 1; Walsh et al., 2023). Segments are combinedwith one anotherusing predictable rules into single discrete calls where the transition between segments within a call is delineated by sudden spectral shifts or a very brief period of silence (<0.01s) (e.g. NLDS Figure 1; Walsh et al., 2023). Some segments may also be produced in isolation as ‘single-segment’ discrete calls: *NL* is a low-level general disturbance alarm call and *DS* and *LH* occasionally occur in isolation though their function, if any, remains unknown (Mason, 2025; Walsh, 2024). Call sequences are single vocal events in which two to 15 single- and multi-segment calls are combined using predictable second-order rules (where each call depends on both preceding calls occurring together; Mason, Walsh, et al., 2026; Figure 1), with brief silences of ≤0.5s between calls (Walsh et al., 2023). While the meaning of every individual call and sequence has not yet been feasible to determine in a repertoire of over 100 call sequences, discrete calls appear to predominantly function as alarm calls (Dutour et al., 2020, 2023) with initial evidence suggesting call sequences refer to more specific contexts such as aerial predator threats or conspecific intruders to the group territory (Mason, 2025; Walsh, 2024).

**Figure 1.**
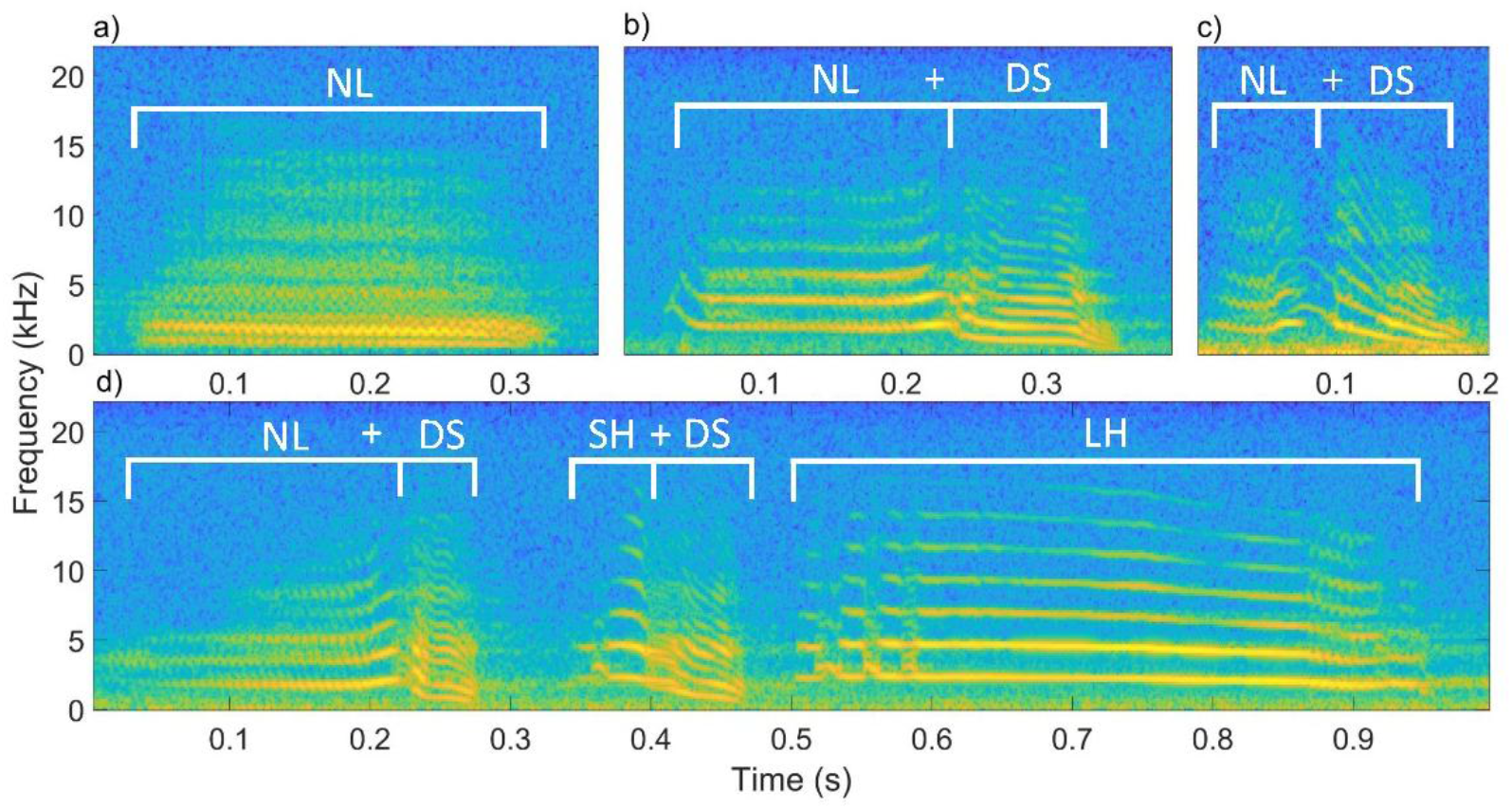
Examples of magpie vocalisations with (a) showing a discrete *NL* call, (b) and (c) two *NLDS* calls where segment duration, as well as the space between segments vary and (d) showing an *NLDS-SHDS-LH* sequence. Segments (the smallest vocal unit) can either be produced in isolation (e.g., *NL*) or can be combined into discrete calls such that segments within the call are distinguishable by sudden spectral shifts or very brief silences of <0.01s (e.g., *NL* + *DS* in *NLDS*). Discrete calls can either be produced in isolation such that they are separated from distinct vocal events by periods of silence >0.5s or combined into call sequences where individual calls are separated by periods of silence ≤0.5s, such as in (d). Segments are labelled according to established spectrographic structure; refer to Walsh et al., 2023 for further detail on magpie vocal classification.

### Data collection

Data collection for this study was conducted in the breeding season of 2022-2023, during the austral summer, and resulted in a final sample size of 11 fledglings from 11 mothers from 8 wild-living magpie groups situated in the metropolitan suburbs of Crawley (31.97°S, 115.82°E; n = 5 groups) and Guildford (31.89°S, 115.97°E; n = 3 groups) in Perth, Western Australia. Fledglings are distinguishable from adults by their lack of sexually mature plumage: fledglings have mottled grey plumage, whereas adult males are black with white backs (males) or with black and white mottled backs (females). Individuals are identifiable through unique, coloured rings or distinctive physical features (e.g. scarring or plumage). SLM followed individuals from their first week as fledglings to 200 days post-fledging, thus capturing both the initial dependent period of development (first 100 days), and the beginning of independence and integration into the broader social group (second 100 days post-fledging) (Ashton et al., 2018). Each fledgling was followed for one hour per week during the first 100 days, and then once every three weeks until day 200. During these focal follows, continuous audio recording captured the vocalisations of the focal fledgling and their group members. Caller identity for each vocal event was noted in real time using voice notes recorded on the same audio track. If the caller could not be identified, it was recorded as ‘unknown’. In cases where it was possible to determine whether a vocalisation came from an adult or a non-focal fledgling (due to known general locations of group members at the time), vocalisations from ‘unknown’ adults could still be included in the analyses. Non-focal fledgling vocalisations were not included as these tended to be individuals whose fledge dates were unknown (so developmental changes could not be assessed) or who died within several weeks (fledgling mortality during the study was 45%). No juveniles—individuals from a previous breeding season who are not yet sexually mature—were present in any groups during the study. See Mason, King et al. (2026) for further details on data collection.

### Audio preparation

Audio recordings were processed in Adobe Audition (v22.3). The start and end of individual vocal events were marked, and each event named in the audio file according to its composition: whether it was a discrete call or a sequence, and by the segments it contained (*sensu* (Walsh et al., 2023)) (Figure 1). The identity of the caller for each vocal event was noted in voice notes on each recording and from this, their ID was added to the marker description (see Mason, King, et al., 2026 for further detail). All vocal events of both fledglings and adult group members were extracted using custom MATLAB scripts before their spectrograms were visually inspected. Vocalisations that were deemed to be of high enough quality for acoustic analysis (based on visual assessment of good signal to noise ratio) were selected and cut into their constituent segments (e.g. *NLDS-SHDS-LHDS* became clips of *NL, DS, SH* and *LH*). This ensured acoustic analysis was assessing the structure of the sounds themselves and was not impacted by variable periods of silence between segments or calls. This also meant where one segment was of poor quality due to noise interference affecting part of the vocal event, the other segment(s) could still be used. Information on whether the segment was cut from a multi-segment call or had been produced as a single segment call (either in isolation or as a single-segment call within a sequence) was retained so that acoustic differences between segments produced alone and those produced in conjunction with another segment in a multi-segment call could be analysed. Information on the original source vocalisation was retained for each segment in preparation for analysis, so that their non-independence to other segments from that vocal event could be identified. However, a maximum of three segments were taken from any single vocal event, and the overall sample size (see below) was large enough that individual vocal events were not leveraging the results. A total of 40 high quality segments collected during a preliminary investigation in 2021 (Mason, 2021) following the same focal protocol as Mason, King et al. (2026) were included from an additional 6 fledglings from that breeding season. This helped to boost the sample size for certain segment types/developmental stages that were underrepresented. Prior to finalising the dataset, we analysed an additional vocalisation previously classified as a ‘grunt’ that magpies produce as a contact call from the nestling stage through to adulthood, strongly suggesting an innate basis (Mason, 2025). From visual inspection, spectrograms of *NL*s appeared to be a graded, higher-arousal version of a grunt. We assessed acoustic similarity in good quality fledgling grunts and *NL*s produced as single-segment discrete calls (i.e., isolated *NL*s) and found that grunts were acoustically indistinct from *NL* (see supplementary material Figure S.1). As such, grunts were relabelled as *NL*s and included as such in the subsequent analyses. The final segment dataset comprised 1681 individual segments (n = 838 adult segments and n = 843 fledgling segments) cut from 1138 vocal events (n = 558 adult vocal events—252 discrete calls, 306 sequences; n = 580 fledgling vocal events—332 discrete calls, 248 sequences). In addition to the segment-level analysis, there was sufficient sample size (n = 180) of one commonly produced multi-segment call—*NLDS* (Figure 1)—to compare whether fledgling productions (n = 113) and adult productions (n = 67) of an entire call (as opposed to the constituent segments) were acoustically distinct.

### Data preparation

For each analysis conducted, a CSV was created containing the individual audio filenames in the rows of the first column, and a predictor variable to be analysed in the second column. To examine the potential for acoustic development over time we analysed between-segments differences of (i) the segments of fledglings and adults together, to assess the presence of overall differences between the two, and (ii) fledgling segments (without adults)—to investigate possible finer scale changes in fledgling production from early to late development. We then assessed within-segment differences in fledglings to shed light on possible segment-specific patterns not apparent when all segment types were included simultaneously. For both between- and within-segment analysis, we assessed if a range of predictor variables (analysed one at a time) significantly correlated with the distribution of data in each of the analyses: (i) segment label (i.e., *DS, NL, LH*, and *SH*), followed by (ii) age group (i.e., fledgling or adult) in the combined adult-fledgling analyses, (iii) fledgling age (grouped into weeks 1-10, 11-20 and 21-30 post-fledging respectively) in the fledgling-only analyses, (iv) group location (Crawley or Guildford—previously shown to cause variation in adult segments, Walsh et al., 2023 ) and (v) group identity (ID). The within-segment analysis could only be conducted on *NL* and *DS* segments due to the lower production of *SH* and *LH* segments in early fledgling development (UMAP requires a minimum of 100 samples, Thomas et al., 2022). Segment source (either ‘cut’ for those cut from multi-segment calls or ‘isolated’ for those produced as a single segment call or as a lone call within a sequence) was included as an additional predictor in the within-segment analysis of *NL* to test for the effects of co-articulation found in adult segments in Walsh et al. (2023). *DS* was not produced in isolation enough to be able to compare cut and isolated *DS*. Finally, we conducted a within-call analysis of fledgling and adult *NLDS* calls simultaneously, including age group, fledgling age, group location and group ID (as above) separately as predictor variables.

Individual identity was not included as a predictor in any of the analyses, as over 20 individuals were included in many of the datasets, creating too many predictor classes to explain UMAP clustering patterns to a meaningful degree in the sample sizes available. Instead, the potential for certain individuals to leverage results was mitigated by balancing the datasets. To balance each of the datasets, any segment class or individual with a sample size more than twice the average of the others was downsampled. In such cases, files were randomly removed from the overrepresented class or individual until their sample size was reduced to no more than twice the average of the remaining classes or individuals. When balancing adult data, many of the individual callers were unknown individuals, so data was balanced according to group identity instead, using the same rule of no more than double the average samples of the other groups (sample sizes given in *Audio preparation* are after down sampling) See Thomas et al. (2022) for detail on data structuring for UMAP analysis.

### Acoustic analysis

The method for acoustic analysis followed Thomas et al., (2022), with necessary adaptations made to better reflect acoustic parameters for magpie vocalisations. All recordings had a bandpass filter applied set at 0.3-8kHz, to encompass the frequency range in which the magpie vocalisations analysed lay. Spectrograms were created via short-time Fourier transformation with a Mel filterbank set to the same range as the bandpass filter, applied to logarithmically scale the spectrograms. Other parameters were kept the same as in Thomas et al (2022). Spectrograms were projected into latent space using UMAP from *umap-learn* (McInnes et al., 2020). Only one of the predictor variables (described in *Data preparation* above) can be analysed at a time in UMAP. Spectrograms for each cluster analysis were plotted as points colour coded by the classes of the predictor variable being investigated, so that correlations between these variables and the distribution of data could be visually assessed.

### Cluster evaluation

The extent to which each predictor described the distribution of data was evaluated by assessing distances between data points belonging to the same predictor class, relative to the nearest data points from another class. This was quantified by the silhouette coefficient (*SIL*), a global metric calculated as the average of each silhouette score (*sil*)—a local measure of how close an *individual* datapoint is to others in its own class compared to those in other classes (Thomas et al., 2022). Averaging these values to obtain *SIL* provides an overall indication of how well the predictor accounts for the clustering of the data. Positive *SIL* values indicate that overall, spectrograms are nearest to spectrograms of the same cluster, while a negative value indicates that spectrograms are closer to those of other clusters.

### Statistical analysis

To assess whether the clustering explained by the predictor is greater than what would be expected by chance (i.e., if the data had no underlying structure related to that predictor), we conducted permutation tests. Specifically, predictor labels were randomly distributed for each analysis across 10,000 permutations, keeping each spectrogram’s original position in latent space but shuffling the predictor labels across datapoints. For each shuffled permutation, *sil* was calculated for each datapoint (yielding 10,000 *sil* scores per spectrogram), and averaged to obtain *SIL* for each permuted data set. An empirical p-value was then calculated by determining the proportion of permuted *SIL* that were equal to or greater than the observed *SIL*, providing a non-parametric measure of significance for how well the predictor explains clustering beyond chance. Observed *SIL* (hereafter obs*SIL*), mean permuted *SIL* (hereafter perm*SIL*), and corresponding p values derived from the permutation tests are presented in the results.

## Results

### Between-segment analysis

#### Adults versus fledglings

When combined, adult and fledgling segments were best explained by the established segment labels (*DS, LH, NL* and *SH*) (obs*SIL* = 0.522; Figure 2a) which explained the distribution of data significantly more than chance expectation (perm*SIL* = -0.030, p <0.001). No difference was found between adult and fledgling spectrograms (obs*SIL* < 0.001, perm*SIL* < 0.001, p = 0.429; Figure 2b).

**Figure 2.**
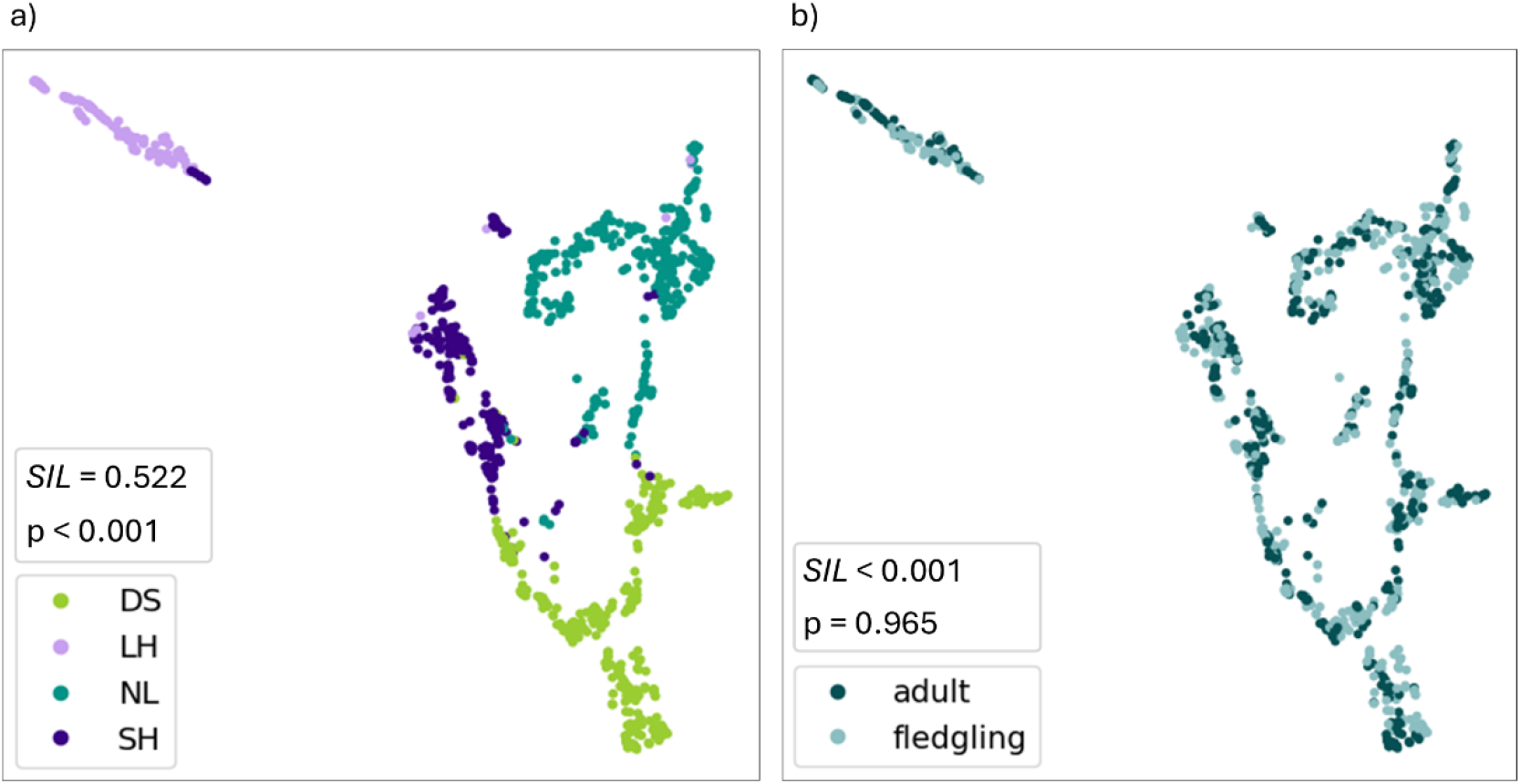
Spectrograms of 4 vocal segment types—’*down sweep’ (DS)* (n = 356), ‘*long high’ (LH)* (n = 141), ‘*noisy line’ (NL)* (n = 356) and ‘*short high’ (SH)* (n = 228)—produced by 20 fledgling magpies aged between 1- and 30-weeks post-fledging (N = 534) and adults from their social environments (N = 547) projected into UMAP latent space. (a) shows the data labelled by segment type, where clusters are well explained by the labels (silhouette coefficient, *SIL* = 0.522), significantly more so than expected by chance (mean permuted *SIL* = -0.030; p <0.001). (b) shows the data labelled by whether the segment was produced by an adult or a fledgling, regardless of segment type. No difference was found between the two age groups, with no distinct clustering of fledgling or adult datapoints (*SIL* = <0.001, mean permuted *SIL* <0.001, p = 0.429).

#### Fledglings

When analysing only fledgling segments, the established segment labels remained the best predictor for explaining the clustering of data (obs*SIL* = 0.466, perm*SIL* = -0.037; p <0.001) (Figure 3a). *LH* segments clustered most distantly from other segment types, but all segment classes were well-separated with minimal overlap (Figure 3a). In contrast, fledgling age did not explain the distribution of data (obs*SIL* = -0.026, perm*SIL* = -0.022; p = 0.733; Figure 3b).

**Figure 3.**
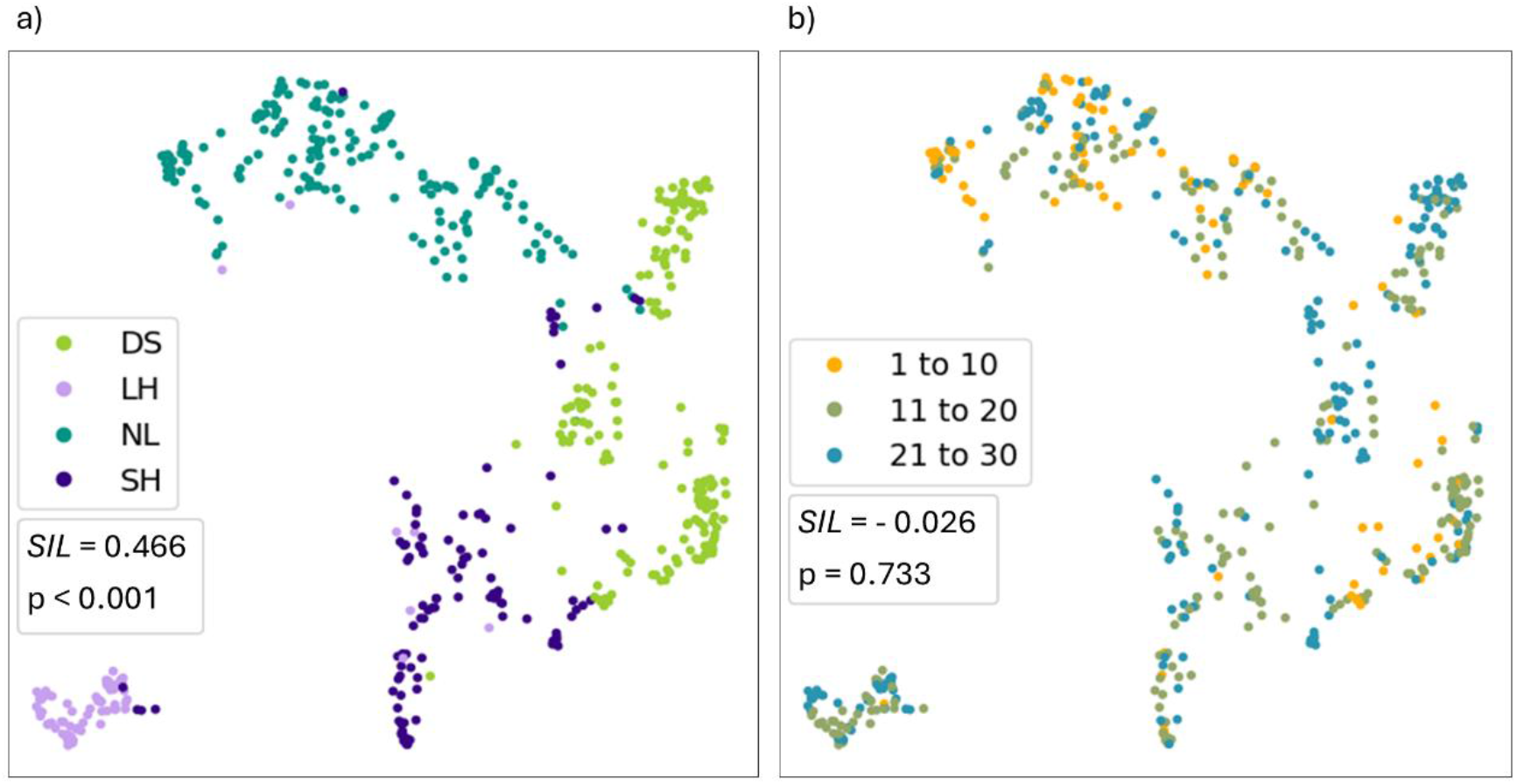
Spectrograms of 4 vocal segment types—*’down sweep’ (DS)* (n = 178), ‘*long high’ (LH)* (n = 73), ‘*noisy line’ (NL)* (n = 178) and ‘*short high’ (SH)* (n = 105)—produced by 20 fledgling magpies aged between 1- and 30-weeks post-fledging (N = 534) projected into UMAP latent space. (a) shows the data labelled by segment type, where clusters are well explained by the labels (silhouette coefficient, *SIL* = 0.466, mean permuted *SIL* = -0.037, p <0.001). (b) shows the data labelled by fledgling age (1 to 10, 11 to 20 and 21 to 30 weeks post-fledging), where clusters are not well explained (*SIL* = -0.026; mean permuted *SIL* = -0.022; p = 0.733).

### Within-segment analysis

#### The fledgling NL segment

Fledgling age did not explain the distribution of *NL* segments well (obs*SIL* = -0.004; Figure 4a). Although the fledgling age explained significantly more clustering structure than expected by chance (perm*SIL* = –0.029; *p* < 0.001), the overall silhouette coefficient remained negative, indicating that the clustering based on this predictor was weak and possibly caused by an underlying confounding variable. Specifically, age and whether the segment was cut from a multi-segment call or produced in isolation (i.e. segment source) were strongly confounded (χ^2^(2) = 123.73, p < 0.001) such that isolated *NL*s were much more prevalent in weeks 1 to 10 and cut *NL*s, in weeks 11-30. This was reflective of the development of fledgling repertoire over time, with *NL* being the first call produced by any individual, followed by *NLDS* later in development (Mason, King, et al., 2026). This was further confirmed by segment source explaining the distribution of spectrograms much better than fledgling age (obs *SIL* = 0.307; perm*SIL* < 0.001, p < 0.001; Figure 4b).

**Figure 4.**
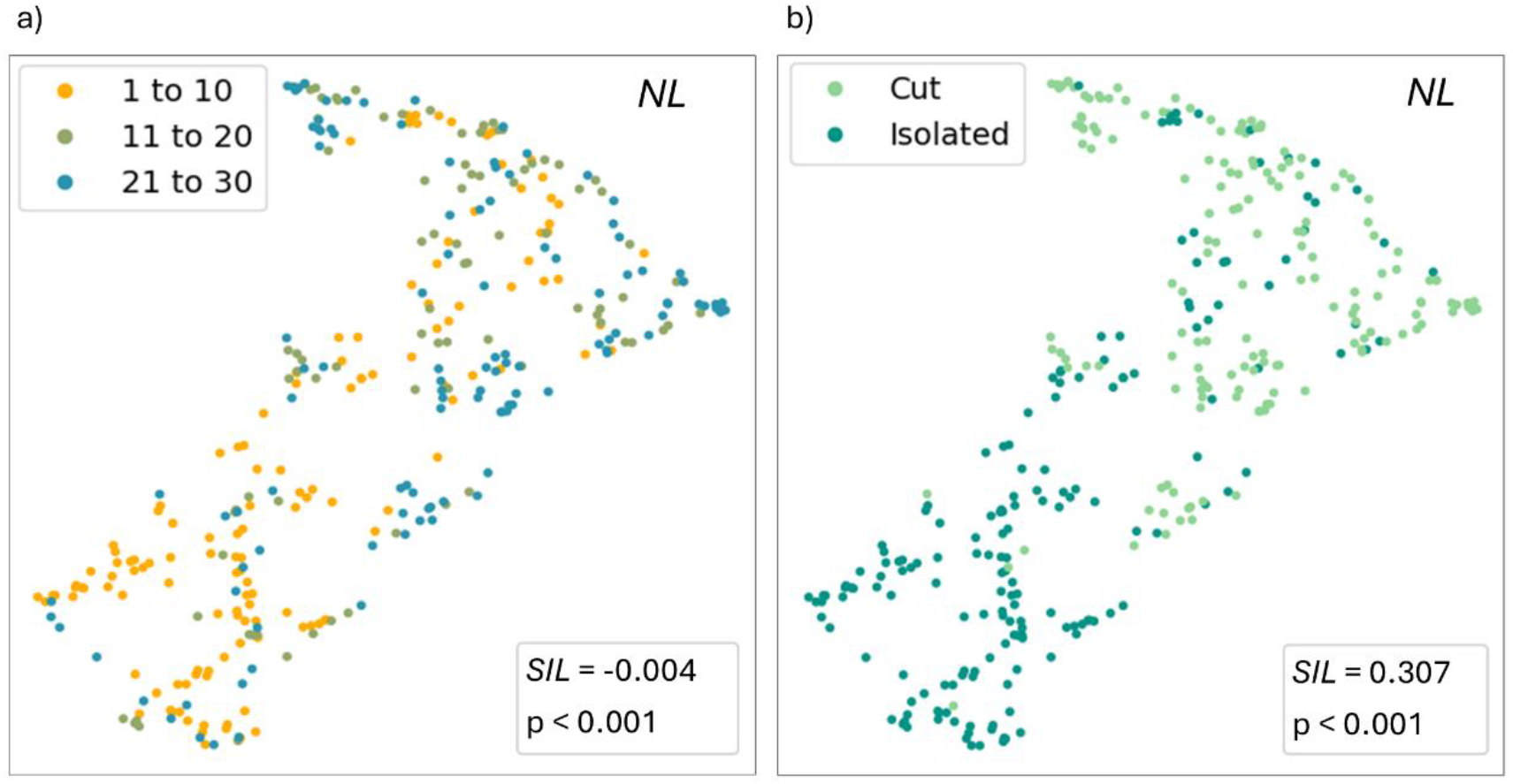
Spectrograms of *‘noisy line’* (*NL*) vocal segments (n = 341), produced by 15 fledgling magpies aged between 1- and 30-weeks post-fledging projected into UMAP latent space. (a) shows the distribution of spectrograms labelled by fledgling age (1 to 10, 11 to 20 and 21 to 30 weeks post - fledging) where the clusters are not well explained (silhouette coefficient *SIL* = -0.004) but the distribution of points under the predictor is significantly different from random (mean permuted *SIL* = –0.029; *p* < 0.001), likely due to confounding of age with segment source. (b) shows the spectrograms labelled by segment source—i.e. whether they were produced in isolation (as a stand-alone call or a single-segment call within a sequence) or were cut from multi-segment calls. Segment source explains the clustering of data well (*SIL* = 0.307, mean permuted *SIL* < 0.001, p < 0.001).

Group location also significantly explained the distribution of *NL*s (obs*SIL* =0.034, perm*SIL* < -0.001, p < 0.001), but again, clusters were not well defined, leading to a weak silhouette coefficient (see supplementary material Figure S.4. for further investigation). Group ID did not explain the distribution of the data (obs*SIL* -0.231, perm*SIL* = -0.154, p = 0.965).

#### The fledgling DS segment

Spectrograms of fledgling *DS* segments did not cluster well according to age groups (obs*SIL* = -0.033, perm*SIL* = -0.040, p = 0.305; Figure 5a). Group location did explain some vocal clustering (obs*SIL* = 0.163) and significantly more than expected by chance, despite a relatively low obs *SIL* (perm*SIL* < - 0.001, p < 0.001; Figure 5b). Group ID did not explain the distribution of data (obs*SIL* = -0.182, perm*SIL* = -0.265, p = 0.051).

**Figure 5.**
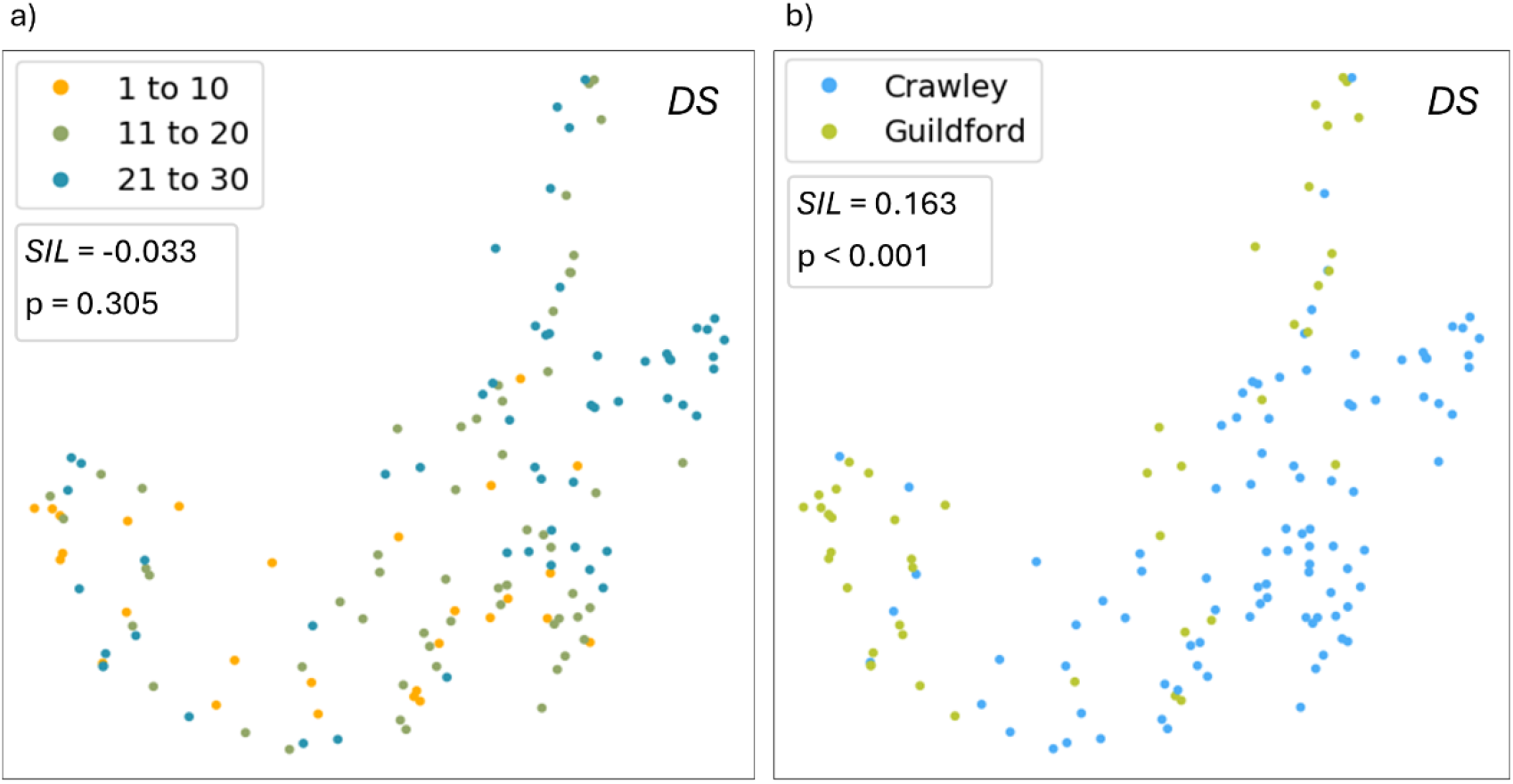
Spectrograms of ‘*down-sweep’* (*DS*) vocal segments (n = 141) produced by 16 fledgling magpies aged between 6- and 30-weeks post-fledging, projected into UMAP latent space. (a) shows the spectrograms labelled by fledgling age (1 to 10, 11 to 20 and 21 to 30 weeks post-fledging) where the clusters are not well explained (silhouette coefficient *SIL* = -0.033, mean permuted *SIL* = -0.040, p = 0.305). (b) shows the spectrograms labelled by whether they were produced by fledglings in the Crawley or Guildford populations, which better explains the clustering of data (*SIL* = 0.163, mean permuted *SIL* < -0.001, p < 0.001).

### Within-call analysis: *NLDS*

We found no difference between the *NLDS* productions of fledglings and adults (obs*SIL* = -0.011, perm*SIL* < -0.001, p = 0.898; Figure 6a) nor when the data was labelled by finer scale age groups (1 to 10, 11 to 20, 21 to 30 week post-fledge and adult) (obs*SIL* = -0.065, perm*SIL* = -0.065, p = 0.541; Figure S.6a.). Instead, group location best explained the distribution of data (obs *SIL* = 0.155, perm*SIL* = - 0.001, p < 0.001; Figure 6b). The effect of group location remained stable even following further down-sampling of the more numerous Crawley datapoints (Figure S.6c.). Group ID explained the distribution of data significantly more than chance but poorly, with a negative obs*SIL* (Figure S.6b), and was likely due to confounding with location effects—which explained the distribution of data better—or individual differences (see supplementary material Figure S.6. for further detail).

**Figure 6.**
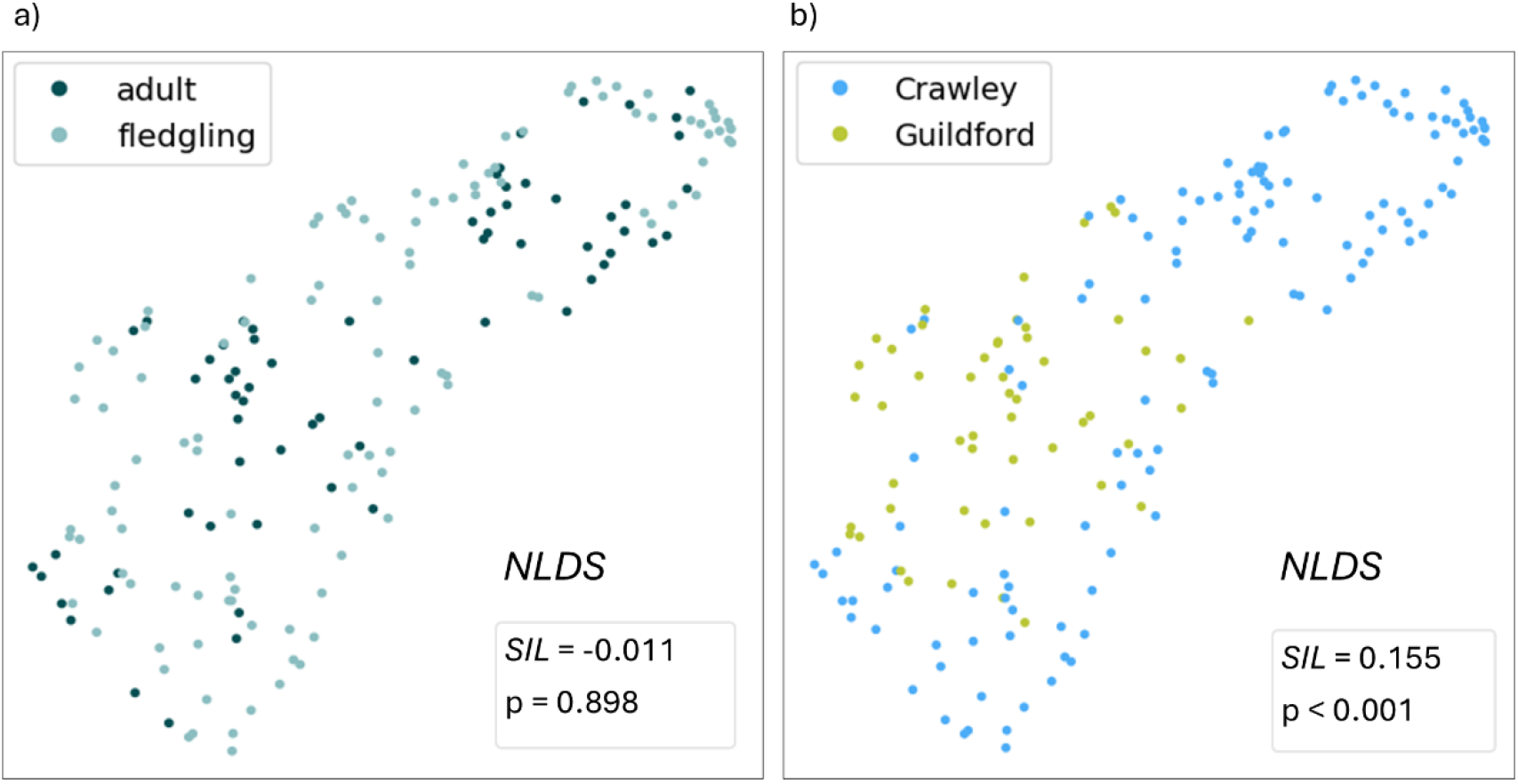
Spectrograms of ‘*noisy line-down sweep’* (*NLDS*) calls (N = 180) produced by 18 fledgling magpies (n = 113) aged between 4- and 30-weeks post-fledgling, and by adults in their social groups (n = 67). (a) shows the spectrograms labelled by whether they were produced by an adult or fledgling, which does not explain the data well (silhouette coefficient *SIL* = -0.011, mean permuted *SIL* < -0.001, p < 0.001). (b) shows the spectrograms labelled by whether they were produced by individuals in the Crawley or Guildford populations, which better explains the clustering of data (*SIL* = 0.163, perm*SIL* = -0.001, p < 0.001).

## Discussion

We set out to determine whether the building blocks of the magpie call sand sequences undergo acoustic development during ontogeny. Mason, King et al. (2026) indicated that, while sequences show evidence of social learning, the vocal elements within them may be innate in origin given their earlier appearance in the repertoire and the lack of social influence on their development. Here, we show that fledgling and adult segments are indistinct regardless of age at production, clustering according to the established segment labels (Walsh et al., 2023). Furthermore, within-segment analyses of fledgling *NL* and *DS* segments revealed coarticulatory effects (in *NL* produced alone versus within a call, Figure 4b) and effects of geographic location (in both *DS*, Figure 5b; and *NL*, Figure S.4a, d and e) previously suggested in adults (Walsh et al., 2023) are already present at the fledging stage. Additionally, we showed that even the multi-segment call, *NLDS*, is acoustically indistinct between fledglings and adults. These findings indicate that the underlying vocal elements comprising the magpie call and call sequence repertoire cannot be learned through vocal production learning. Together with the evidence of social learning in Mason, King et al. (2026), these results provide support for a developmental trajectory in which magpies learn how to correctly combine pre-existing, innate calls into complex sequences from their social contacts.

Given the recency with which semantic call sequences have been discovered in animal systems, there has been little investigation into the ontogenetic processes through which call sequences emerge. Both vocal production learning and vocal usage learning—learning from experience a new way in which to use an existing signal (Janik & Slater, 2000; Vernes et al., 2021)—are well documented in shaping the sequences of syllables within bird song (Beecher & Akçay, 2020; Hultsch & Todt, 1989; Marler & Peters, 1977; Thorpe, 1958), but the role of learning in semantic call sequences remains poorly studied outside of our work on magpies (Mason, 2025; Mason, King, et al., 2026; Mason, Walsh, et al., 2026). Innate and learned vocalisations are almost always discussed as antithetical mechanisms, but as our understanding of the neurobiological processes behind vocal learning grows, it becomes increasingly apparent that learned vocalisations are often the product of underlying innate building blocks (Mooney, 2020; Rose et al., 2022; Vernes et al., 2021). In primates for example, the innate basis of calls is well established (Janik & Knörnschild, 2021), but variation in the call order of a two-call sequence produced by two chimpanzee populations was attributed to different social dynamics in the two groups and suggested that, despite the apparent innate basis of the individual calls, the order in which they are produced may be acquired through vocal usage learning earlier in life (Girard-Buttoz, Bortolato, et al., 2022). Additionally, a study by Bortolato et al. (2023) revealed a protracted period of vocal development in chimpanzees of ∼10 years, with increases in sequence length coinciding with key social milestones during development, though no direct evidence of learning was shown. These studies show the capacity for call sequencing in chimpanzees is socially driven, even if the vocal building blocks are genetically determined.

Magpies, unlike primates, are capable of vocal production learning, and yet our results suggest magpie segments are innate. While we would need to isolate magpie fledglings from as early as brooding (see Kleindorfer et al., 2024 for evidence of possible in-ovo learning) to conclusively show their segments are innate, the evidence provided here, together with the fact that calls emerge early in development, showing little change with age and no effect of social environment (Mason, King, et al., 2026), strongly suggests that all fledglings start life with the same vocal building blocks (i.e. segments and possibly multi-segment calls). In a concurrent study we showed that, like adults, fledglings use second-order rules to structure their sequences (Mason, Walsh, et al., 2026). However, fledgling sequence structure is far less stable, producing many novel transitions infrequently, suggesting they are vocal practice rather than stable fledgling-specific sequences (Mason, Walsh, et al., 2026). Together, the fact that the same calls are combined into group-specific sequence repertoires that fledglings learn from social contacts (Mason, King, et al., 2026) and that the rules of combining calls develop over time (Mason, Walsh, et al., 2026) but not the acoustic quality of the comprising segments, strongly suggest vocal usage learning, and not production learning underpins call sequences.

The lack of acoustic development apparent in magpie segments—while allowing us to support their basis as innate vocalisations—is surprising. Even species with genetically determined repertoires have been observed to undergo a period of vocal development due to maturation of the vocal tract (Fitch & Hauser, 1995; Seyfarth & Cheney, 1997), changes in lung capacity (Hammerschmidt et al., 2001), or simply practice (Seyfarth & Cheney, 1997; Snowdon et al., 1997) (though see Takahashi et al., 2015, 2017 for evidence of social vocal feedback in infant marmosets). One possible explanation for the lack of acoustic development apparent here, is that the staggered emergence of vocal segments provides sufficient practice for each subsequent segment to be produced in adult-like form. Segments were not equally represented in each age class—*NL* was the only segment produced from week 1 and produced reliably as an isolated call before week 10, comprising 65% of the segments in the week 1 -10 category. While isolated *NL*s and fledgling grunts were found to be indistinct (Figure S.1.), they were initially labelled as separate vocalisations based on perceivable differences in arousal state and intensity during production. Many animals produce innate care-eliciting vocalisations from the very start of life—such as cries in human newborns (Eibl-Eibesfeldt, 1973) and begging in many bird species (including magpies) (Leonard & Horn, 2001). Similarly, magpie grunts appear to emerge immediately, helping nestlings maintain vocal contact with caregivers. Around the time of fledging, these grunts seem to gradually develop into more intense alarm calls (*NL*s), potentially enabling fledglings to elicit higher arousal and vigilance from caregivers, reflecting their increased vulnerability outside the nest. The period spent grunting in the nest may thus serve as indirect acoustic development of the *NL* segment, leading to their adult-like production as early as week one post-fledging—like the development of adult calls from nestling and fledgling precursors in zebra finches (Taeniopygia spp., Zann, 1996). In contrast, *DS, SH*, and *LH* segments emerge later—at the earliest, in weeks three, seven, and eight, respectively (Mason, King, et al., 2026). Unlike *NL*s, these later-emerging segments may lack equivalent precursors during the nestling phase—possibly because they do not bear meaning on their own in the same way *NL* does. Instead, the emergence and repeated production of the *NL* in early weeks, along with natural physiological growth may facilitate the development of the vocal tract leading to their seemingly adult quality from first production when they later emerge.

The *NLDS* call—produced by all groups and fledglings (Mason, King, et al., 2026)—and its constituent segments, shows no evidence of acoustic development. Despite this developmental stability, we found significant, albeit weak, differences between Crawley and Guildford populations in *NL* and *DS* segments and their combined *NLDS* call. These differences were consistent across age groups, suggesting either that there is a genetic basis to these population-level acoustic differences or that some degree of vocal production learning occurs earlier in development than we were able to capture here to alter the innate segments into population-specific variants. Variation in the acoustic properties of *NL*s produced in isolation versus those cut from *NLDS* calls reflects coarticulatory effects previously shown in adults (Walsh et al., 2023). The change in the sound of *NL* when it precedes *DS* may also point to a limited form of vocal production learning that alters the underlying innate sound (Recasens, 1999; Vernes et al., 2021). These results highlight the value of avoiding strictly categorising vocalisations as innate *or* learned and prompts further investigation into interactions between innate building blocks and the different forms of learning—production and usage learning—that may build upon them to produce complex vocal systems.

Interestingly, although no differences were found between fledgling and adult productions of segments and *NLDS* calls in this analysis, fledgling call sequences sound perceivably different from those of adults, suggesting there may well be differences in the acoustic structure of whole sequences. For example, the silences between calls within sequences may be longer or more variable in fledglings due to them transitioning between calls less fluidly as they learn to do so, resulting in a less proficient overall sound. Our results indicate fledglings already transition smoothly between segments in early development, producing *NLDS* calls that are acoustically indistinct from those of adults. Since transitioning rapidly between segments within a call likely places greater demands on motor coordination than transitioning between calls during the longer pauses within sequences, these findings suggest that any limitations fledglings face at the sequence level stem from developing cognitive abilities, such as working memory, rather than from motor constraints. In line with this, our concurrent structural study revealed that fledglings produce more variable call orders than adults, supporting that it is not motor constraints of combining calls, but rather the understanding and memory of the correct way in which to do so that limits fledglings from producing adult-like sequences (Mason, Walsh, et al., 2026). Future work could attempt to boost the number of high-quality recordings of common sequence types, as well as other multi-segment calls, and perform UMAP on them as whole vocalisations to compare adult and fledgling acoustic properties at a broader scale. Such work could clarify the presence or indeed absence of acoustic development at the call and sequence level.

## Conclusion

Here, we provide evidence that the building blocks underlying call sequences may be innate. This builds on accumulating evidence that it is the process of combining calls, and not the calls themselves, that is shaped by social experience (Mason, King, et al., 2026; Mason, Walsh, et al., 2026). While it was previously unclear whether combinatoriality in an open-ended vocal learner could resemble that seen in non-human primates, or inform existing evolutionary theories of syntax, these findings—particularly the group-specific usage of shared vocal segments—suggest that usage learning, rather than production learning, may underlie magpie call sequences too. Although the exact neural mechanisms behind these vocalisations remain unclear, these results offer insight into the possible evolutionary drivers of syntactic-like vocalisations. That a species capable of both open-ended vocal production learning and combinatoriality appears to favour combining innate signals rather than learn new ones, suggests that other constraints on communicative capacity—such as working memory limitations (Nowak et al., 2000), or the need for stable shared signal meaning among social groups—may have driven the emergence of combinatoriality. This challenges the primate-centric view that syntax evolved solely to overcome genetically fixed repertoires (Bortolato et al., 2023; Nowak et al., 2000). Instead, our findings support a more flexible account: that syntax evolved convergently as a solution to a variety of challenges associated with expanding communicative capacity.

