## Supplementary material for "Innate building blocks underly socially learned call sequences"

*Grunts versus NL*

The distribution of spectrograms was not significantly different from random (perm*SIL* < -0.001, p = 0.251), suggesting isolated *NL* segments and grunts are indistinct (obs*SIL* = 0.007; Figure S.1.). For all analyses (and the segment counts given in *Methods*), fledgling grunts were relabelled as *NL*s.

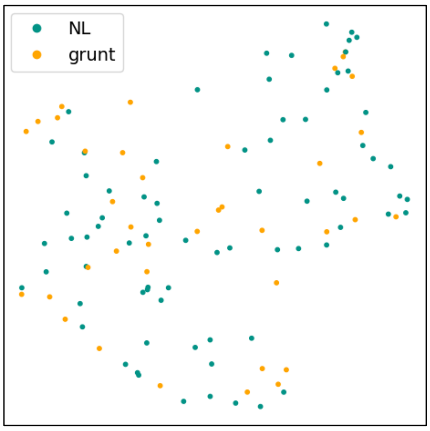


**Figure S.1.** Spectrograms of noisy line (*NL*) (n = 75) and grunt (n = 38) vocalisations of 9 fledgling magpies aged between 1- and 30-weeks post-fledging (N=113) projected into UMAP latent space. The distribution of spectrograms was no different to chance when compared to the average of 10,000 randomly distributed permutations of the data (mean silhouette coefficient of permuted data, perm*SIL* < -0.001, p = 0.251), suggesting they are indistinct (observed silhouette coefficient obs*SIL* = 0.007).

*Between-segment analysis: Adults versus fledglings*

Neither group location (Figure S.2a) nor group identity (Figure S.2b) explained the distribution of adult and fledgling segments (group location: obs*SIL* = -0.003; and group identity; obs*SIL* = -0.213) or was different from random (group location: perm*SIL* = < -0.001, p = 0.642; and group identity: perm*SIL* = -0.167, p = 0.727).

*Between-segment analysis: fledglings*

Neither group location (Figure S.3a) nor group identity (Figure S.3b) explained the distribution of fledgling segments (group location: obs*SIL* = -0.005; and group identity; obs*SIL* = -0.207) or was different from random (group location: perm*SIL* = < -0.001, p = 0.689; and group identity: perm*SIL* = -0.170, p = 0.752).

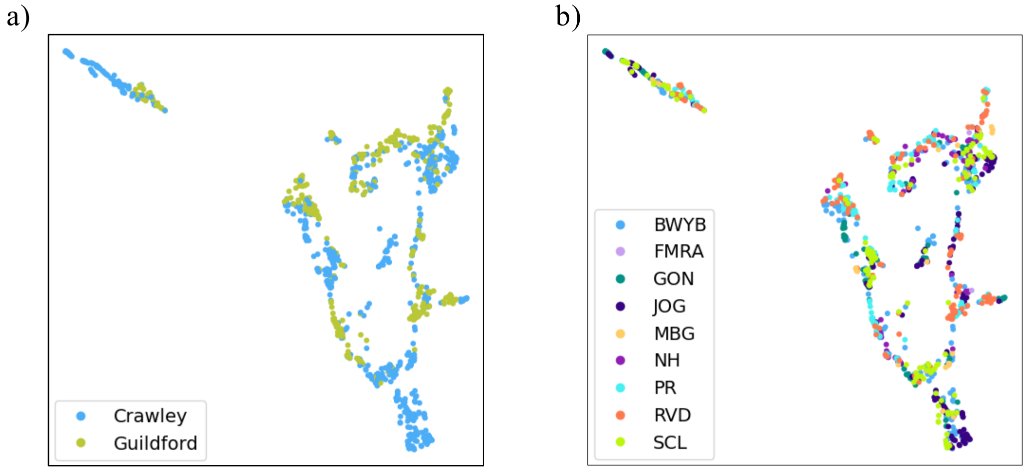


**Figure S.2.** Spectrograms of 4 vocal segment types—*'down sweep' (DS)* (n = 356), '*long high' (LH)* (n = 141), '*noisy line' (NL)* (n = 356) and '*short high' (SH)* (n = 228)—produced by 20 fledgling magpies aged between 1- and 30-weeks post-fledging (N = 534) and adults from their social environments (N = 547) projected into UMAP latent space. (a) shows the data labelled by group location (Crawley or Guildford), where clusters are not well explained by the labels (silhouette coefficient, obs*SIL* = -0.003), are no different to random (mean permuted *SIL,* perm*SIL* < -0.001; p = 0.642). (b) shows the data labelled by group identity of the caller where the distribution of data is not well explained and is no different to random (obs*SIL* = -0.213, perm*SIL* = -0.167, p = 0.727).

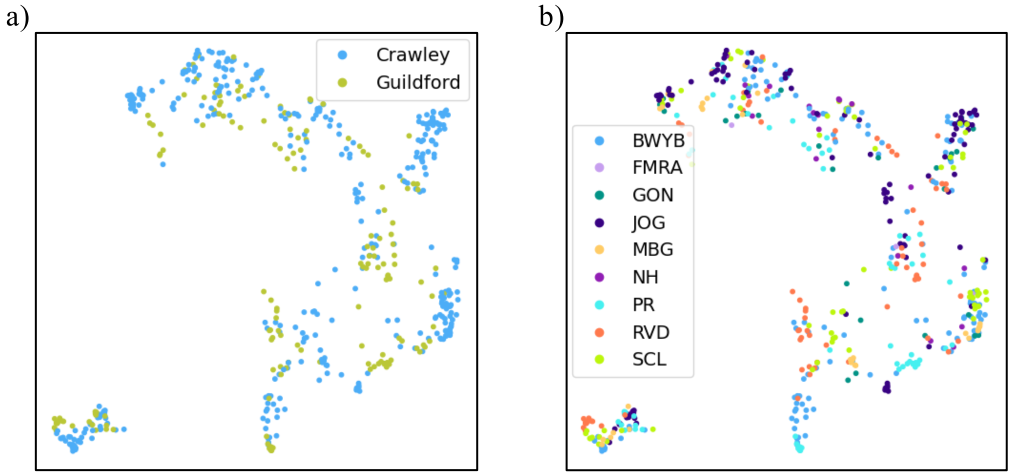


**Figure S.3.** Spectrograms of 4 vocal segment types—*'down sweep' (DS)* (n = 178), '*long high' (LH)* (n = 73), '*noisy line' (NL)* (n = 178) and '*short high' (SH)* (n = 105)—produced by 20 fledgling magpies aged between 1- and 30-weeks post-fledging (N = 534) projected into UMAP latent space. (a) shows the data labelled by group location (Crawley or Guildford), where clusters are not well explained by the labels (silhouette coefficient, obs*SIL* = -0.005), are no different to random (mean permuted *SIL,* perm*SIL* < -0.001; p = 0.689). (b) shows the data labelled by group identity of the caller where the distribution of data is not well explained and is no different to random (obs*SIL* = -0.207, perm*SIL* = -0.170, p = 0.752).

*Within-segment analysis: NL*

When labelled by group location, the distribution of fledgling *NL* segments was found to be poorly explained, but still significantly different from chance (obs*SIL* = 0.034, perm*SIL* < -0.001, p value < 0.001; Figure S.4a). Group identity did not explain the data well, nor was it significantly different from chance (obs*SIL* = -0.231, perm*SIL* = -0.154, p = 0.965; Figure S.4b). Given segment source was found to explain the distribution of fledgling *NL*s best (Figure 5.b), we relabelled the data by both group location and segment source to see if there was an interaction between the two that was causing the significance found with group location, but this did not explain the data well and was insignificantly different from chance (obs*SIL* = -0.0412, perm*SIL* = -0.049, p-value = 0.257; Figure S.4c). To further investigate the significant albeit weak impact of group location on the acoustic structure of *NL*s, we separated the data set by segment source and analysed group location as a predictor once more. In both cut (Figure S.4d) and isolated *NL*s (Figure S.4e), group location was a similarly weak but significant predictor of the distribution of data (in cut *NL*s: obs*SIL* = 0.086, perm*SIL* < -0.001, p < 0.001; and isolated *NL*s: obs*SIL* = 0.045, perm*SIL* < 0.001, p = 0.001). However, given that spectrograms from both locations were spread across the distribution of data with no clear clustering—with obs*SIL* remaining low throughout—and that group location was not previously shown to explain the distribution of adult *NL*s (Walsh et al., 2023), it is unclear whether this effect represents a genuine but weak location effect, or simply the effect of slight individual differences nested within location. The latter could not be investigated as a predictor in any of the analyses here due to there being too many individuals to allow this to be a meaningful label in the analysis, and too few samples from individual fledglings to run the analysis with a smaller number of individuals.

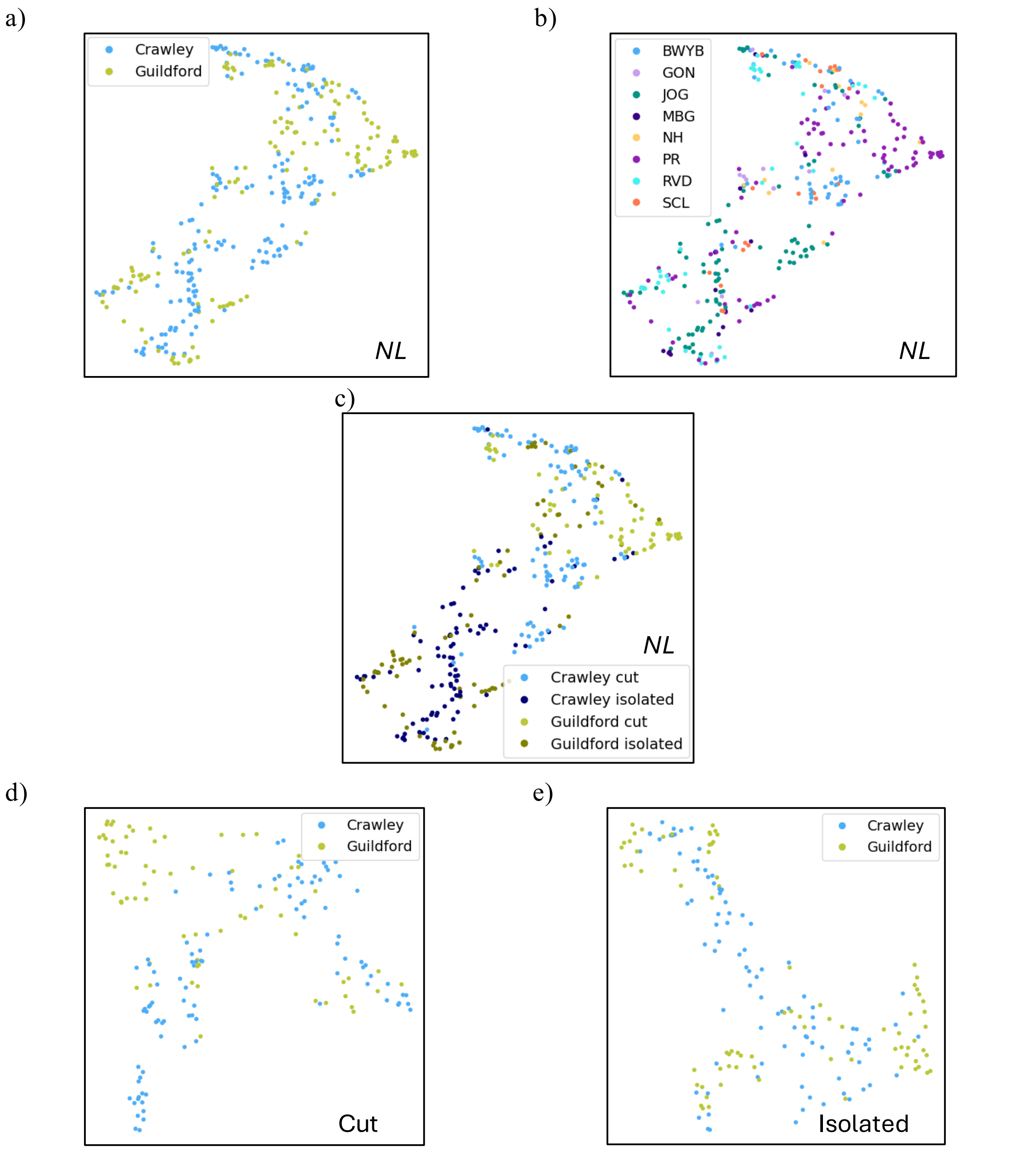


**Figure S.4** Spectrograms of *'noisy line'* (*NL*) vocal segments (including grunts—determined to be the indistinct; Figure S.1.) (n = 341), produced by 15 fledgling magpies aged between 1- and 30-weeks post-fledging projected into UMAP latent space. (a) shows the data labelled by group location (Crawley or Guildford), where clusters are not well explained by the label (silhouette coefficient, obs*SIL* = 0.034) but were significantly different to random (mean permuted *SIL*, perm*SIL* < -0.001; p < 0.001). (b) shows the data labelled by group identity of the caller where the distribution of data is not well explained and is no different to random (obs*SIL* = -0.231, perm*SIL* = -0.154, p = 0.965). (c) shows the data labelled by group location and segment source (cut from a multi-segment call versus produced as a lone segment) where data was not well explained or different from random (obs*SIL* = -0.0412, perm*SIL* = -0.049, p-value = 0.257). (d) and (e) show the data split into cut (n = 167) and isolated (n = 174) segments respectively and labelled by group location, where the distribution of data remains significantly different to random (cut: perm *SIL* < -0.001, p < 0.001 and isolated: perm*SIL* <0.001, p = 0.001) but is still poorly explained by group location (cut: obs*SIL* = 0.086 and isolated: obs*SIL* = 0.045).

*Within-segment analysis: DS*

*DS* was not tested for coarticulatory effects using the segment source variable as they are very rarely produced in isolation. Group ID did not explain the distribution of *DS* segments nor was it significantly different from random (obs*SIL* = -0.182, perm*SIL* = -0.265, p = 0.051; Figure S.5.).

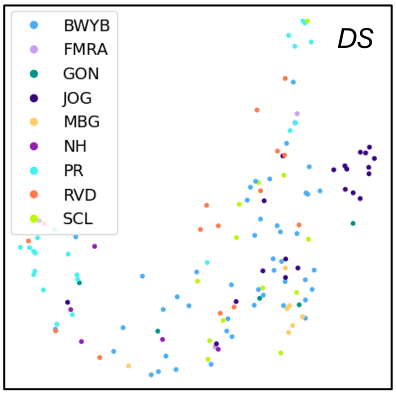


**Figure S.5.** Spectrograms of '*down-sweep'* (*DS*) vocal segments (n = 141) produced by 16 fledgling magpies aged between 6- and 30-weeks post-fledging, projected into UMAP latent space and labelled by group identity. Group ID did not explain the distribution of data well (silhouette coefficient *SIL* = -0.182) nor was it significantly different to random (mean permuted *SIL* = -0.265, p = 0.051).

*Within-call analysis: NLDS*

To confirm the absence of finer-scale acoustic changes to *NLDS* across development, *NLDS* spectrograms were labelled by age group (1 to 10, 11 to 20 and 21 to 30 weeks post fledge or adult; Figure S.6a). This did not explain the distribution of data (obs*SIL* = -0.065) and was not significantly different from random chance (perm*SIL* = -0.065, p = 0.541), further supporting the lack of acoustic changes with age. Group location was the best predictor of *NLDS* (Figure 7). On visual inspection, it looked as though the larger sample size of Crawley spectrograms may be leveraging the result, so to confirm, I reduced the number of Crawley samples (by 24 samples) to further balance the dataset (Figure S.6c). I found the effect of location to be stable following this reduction (obs*SIL* = 0.158, perm*SIL* = <0.001, p < 0.001) suggesting a genuine effect of location. Group ID did not explain *NLDS* data well (obs*SIL* = -0.231) but was significantly different from chance (perm*SIL* = -0.360, p = 0.002), likely due to confounding with group location and the effect found there. Individual differences in production may also lead to some clustering, exaggerated by the greater presence of certain individuals in the dataset (where data could not be perfectly balanced). The true extent of location and individual effects should be clarified further with larger individual sample sizes in future.

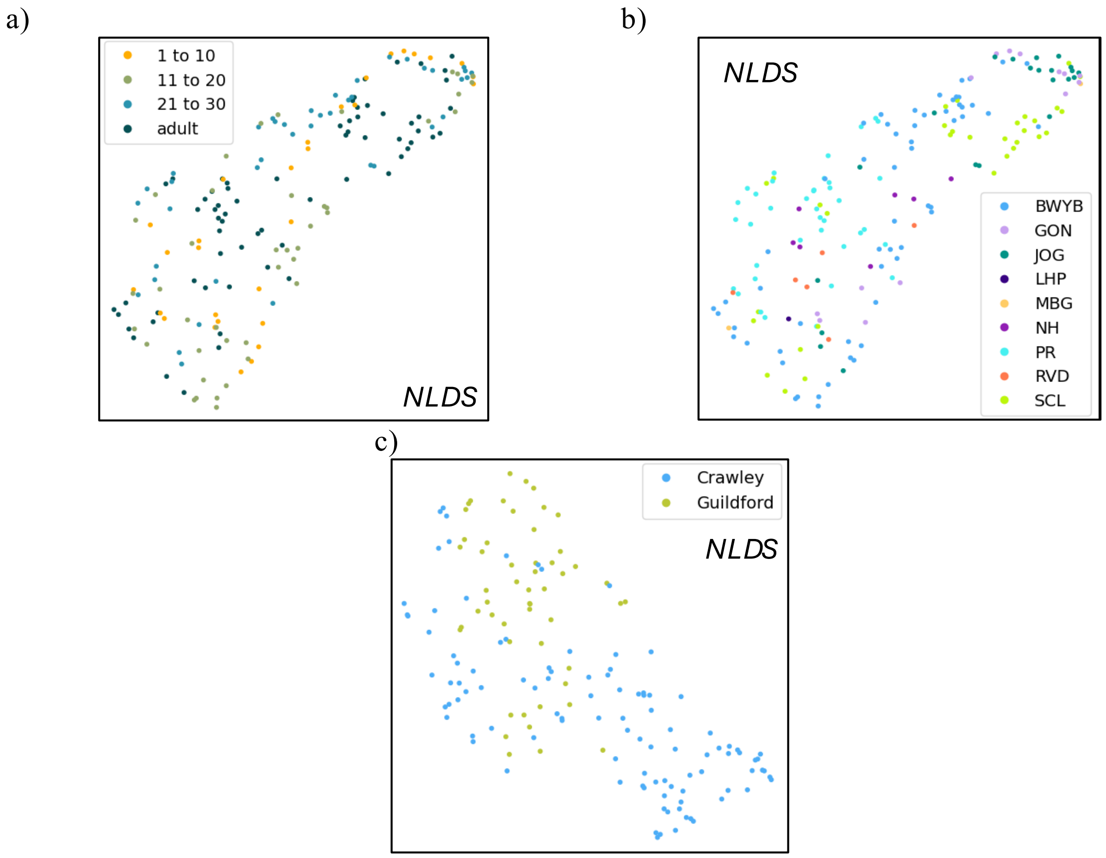


**Figure S.6.** Spectrograms of '*noisy-line—down-sweep'* (*NLDS*) calls (N = 180) produced by 18 fledgling magpies (n = 113) aged between 4- and 30-weeks post-fledgling, and by adults in their social groups (n = 67). (a) shows the data labelled by age group (1 to 10, 11 to 20 and 21 to 30 weeks post-fledging or adult) which did not explain the distribution of data well (silhouette coefficient, obs*SIL* = -0.065) and was not significantly different from random chance (mean permuted *SIL*, perm*SIL* = -0.065, p = 0.541). (b) shows the data labelled by group identity where the distribution of data was not well explained (*SIL* = -0.231) but was significantly different from chance (perm*SIL* = -0.360, p = 0.002), likely on account of weak individual differences exaggerated by greater representation of certain individuals in the dataset. (c) shows a reduced data set (N = 156) where Crawley spectrograms were further downsampled to confirm the presence of a location effect. The effect of group location found in Figure 7 is shown to be stable in this reduced data set (obs*SIL* = 0.158, perm*SIL* < 0.001, p < 0.001).
